# Modeling neurodegenerative diseases in *Drosophila* is conditioned by stress resistance and gut microbiome composition of the reference line

**DOI:** 10.1101/2025.09.03.673979

**Authors:** Xiaojing Yue, Amina Dulac, Amélie Hu, Zohra Rahmani, Serge Birman

## Abstract

*Drosophila* is widely used to study the pathological mechanisms of human diseases *in vivo*, including metabolic and neurological disorders. In these models, disease-induced alterations in locomotion and stress resistance are generally monitored in comparison to healthy control flies, such the white-eyed strain *w*^1118^, used as a reference for normal physiology and behavior. Here we compared two independent *w*^1118^ lines and found that they differed strikingly in their susceptibility to oxidative stress and nutrient starvation, and less markedly in their locomotor performance. Interestingly, modulating the gut microbiome by rearing these flies under axenic conditions increased oxidative stress resistance of the more susceptible, but not the more resistant line, while it had no effect on starvation resistance for both lines. We also found that the stress-sensitive line had higher levels of *Clostridiales* bacteria and of the intracellular endosymbiont *Wolbachia* in the gut microbiota, as well as lower expression levels of immune effectors (antimicrobial peptides and lysozymes) in the head and gut. Both lines nevertheless showed similar susceptibility to pathogenic bacterial infections. In a transgenic Parkinson’s disease model, the stress-resistant background strongly attenuated the progressive locomotor defects induced by pan-neuronal expression of human mutant α-synuclein, but intriguingly not when α-synuclein expression was restricted to a subset of brain dopaminergic neurons in the protocerebral anterior medial (PAM) cluster. These results suggest that taking into account unapparent features of the reference lines could improve the reproducibility and consistency of neurodegenerative disease models in *Drosophila*.

## Introduction

In spite of the large evolutionary distance between humans and insects, *Drosophila* is currently one of the major model organisms to study human diseases and progress in their treatments. This is made possible by the conservation of major transduction pathways and the fact that more than two-third of the human disease-associated genes have orthologues in *Drosophila*, plus the similarity of many internal organs and the ever-increasing molecular tools to study gene expression and function in this genetically tractable animal (Pandey and Nichols, 2011; Perrimon et al., 2016; Pick, 2017; Link and Bellen, 2020). A common way to model hereditary diseases in this organism is to remove or mutate the gene homologous to the altered human gene and monitor the consequences on fly development, physiology and behavior (Ugur et al., 2016). Alternatively, the disease-associated form of human genes can be overexpressed transgenically in the fly or used to replace its *Drosophila* orthologue (Her et al., 2024). Sporadic disorders can be also studied in flies by exposure to drugs or chemicals that are known to be risk factors of human diseases (Rand et al., 2023).

Parkinson disease (PD) is a neurodegenerative disorder that was first modeled in *Drosophila* by pan-neuronal overexpression of human wild-type or mutant α-synuclein, a synaptic protein of poorly understood function that plays a central role in this disease (Feany and Bender, 2000; Auluck et al., 2002; Barone et al., 2011; Riemensperger et al., 2013; Wang et al., 2015; Issa et al., 2018; Mohite et al., 2018; Ordonez et al., 2018; Arsac et al., 2021; Bridi et al., 2021; Girard et al., 2021; Rahmani et al., 2022). This disease can be also modeled in flies by exposure to sublethal doses of pro-oxidant pesticides such as rotenone or paraquat, which are known environmental risk factors for PD (Coulom and Birman, 2004; Chaudhuri et al., 2007; Hosamani and Muralidhara, 2013; Cassar et al., 2015; Shukla et al., 2016; Stephano et al., 2018; Maitra et al., 2019; Rahmani et al., 2022). These models are characterized by age-dependent progressive locomotion defects, increased levels of oxidative stress and loss of dopaminergic neurons in the fly brain. However, these observations were sometimes difficult to reproduce, resulting in negative results which have been more rarely reported (Pesah et al., 2005; Navarro et al., 2014).

The disease development or progression in these models is generally assessed by comparison with healthy control flies, which are used as a reference for normal physiology or behavior. Genetically modified lines used in *Drosophila* studies have been often developed in the *w*^1118^ mutant background, which carries a null mutation in a gene (*white*) encoding an ABC-type guanine transporter involved in eye pigmentation (Hazelrigg et al., 1984). The *w*^1118^ mutant strain was therefore used as a reference in these models, assuming that whatever their origin, various lines of these flies would show comparable wild-type-like performance in relevant assays such as locomotor ability and stress resistance. However, it has been reported that different *Drosophila* wild-type strains (such as Canton-S, Oregon R and others) display variations in age-dependent behaviors (Grandison et al., 2009; Qiu et al., 2017; Zakharenko et al., 2024), and different lines of the same Canton-S wild-type strain can also show differential performance in specific behavior tests (Colomb and Brembs, 2014).

Here we compared two apparently identical but independent lines of the *w*^1118^ strain and observed that they present striking dissimilarities. We found that these two lines, although raised under the same conditions in our laboratory, showed highly divergent susceptibility to oxidative stress and nutrient starvation, and significant differences in locomotion behavior, indicating that these lines are far from being equivalent for use as a reference in neurological disease modelling. Interestingly, rearing the less resistant *w*^1118^ line in germ-free condition somewhat improved its oxidative stress tolerance, and the progressive locomotor defects induced by expression of mutant α-synuclein in all neurons were strongly alleviated in the background of the more resistant *w*^1118^ line. This prompted us to characterize more precisely endogenous features of these control lines that may explain their divergence, focusing on the gut microbiota and innate immune factors such as the anti-microbial peptides (AMPs), which are known to be involved in neurodegenerative processes in *Drosophila* (Kitani-Morii et al., 2021; Lee et al., 2024).

## Results

### Two apparently identical *w*^1118^ lines markedly differ in their oxidative stress and starvation resilience

Critical readout parameters in studies of neurodegenerative disease models in *Drosophila* include susceptibility to oxidative stress and age-related locomotor decline. This is generally assessed by comparing to non-diseased control flies, assuming that there would be no big difference whatsoever between two similar control or wild-type lines (or sub-strains) for those parameters. In order to test this hypothesis, we compared the oxidative stress and starvation resistance of two independent lines of the *w*^1118^ strain available in our laboratory, here named for convenience *w*^lineA^ and *w*^lineB^.

Acute oxidative stress was induced by ingestion of paraquat, a pro-oxidant herbicide commonly used to model drug-induced PD in *Drosophila* (Chaudhuri et al., 2007; Cassar et al., 2015; Shukla et al., 2016; Maitra et al., 2019; Rahmani et al., 2022). Unexpectedly, under strictly similar intoxication conditions, we observed dramatic differences in oxidative stress tolerance between the two lines, with a median survival time (at which point 50% of the flies are dead) of about 19 hours for *w*^lineA^ against 51 hours (2.7 times longer) for *w*^lineB^. Moreover, no *w*^lineA^ fly survived longer than 48 hours, while a few *w*^lineB^ flies were still alive after 96 hours of paraquat exposure (Fig.1A). Furthermore, we found that *w*^lineB^ flies were also much more resistant to nutrient starvation than *w*^lineA^ flies, with a median survival time of about 37 hours for *w*^lineA^ against 60 hours (1.6 times longer) for *w*^lineB^, and a maximal survival time of 59 and 91 hours for *w*^lineA^ and *w*^lineB^, respectively (Fig. 1B). We therefore wondered what could be at the origin of such large variations in stress resistance between two apparently identical control lines.

**Figure 1.**
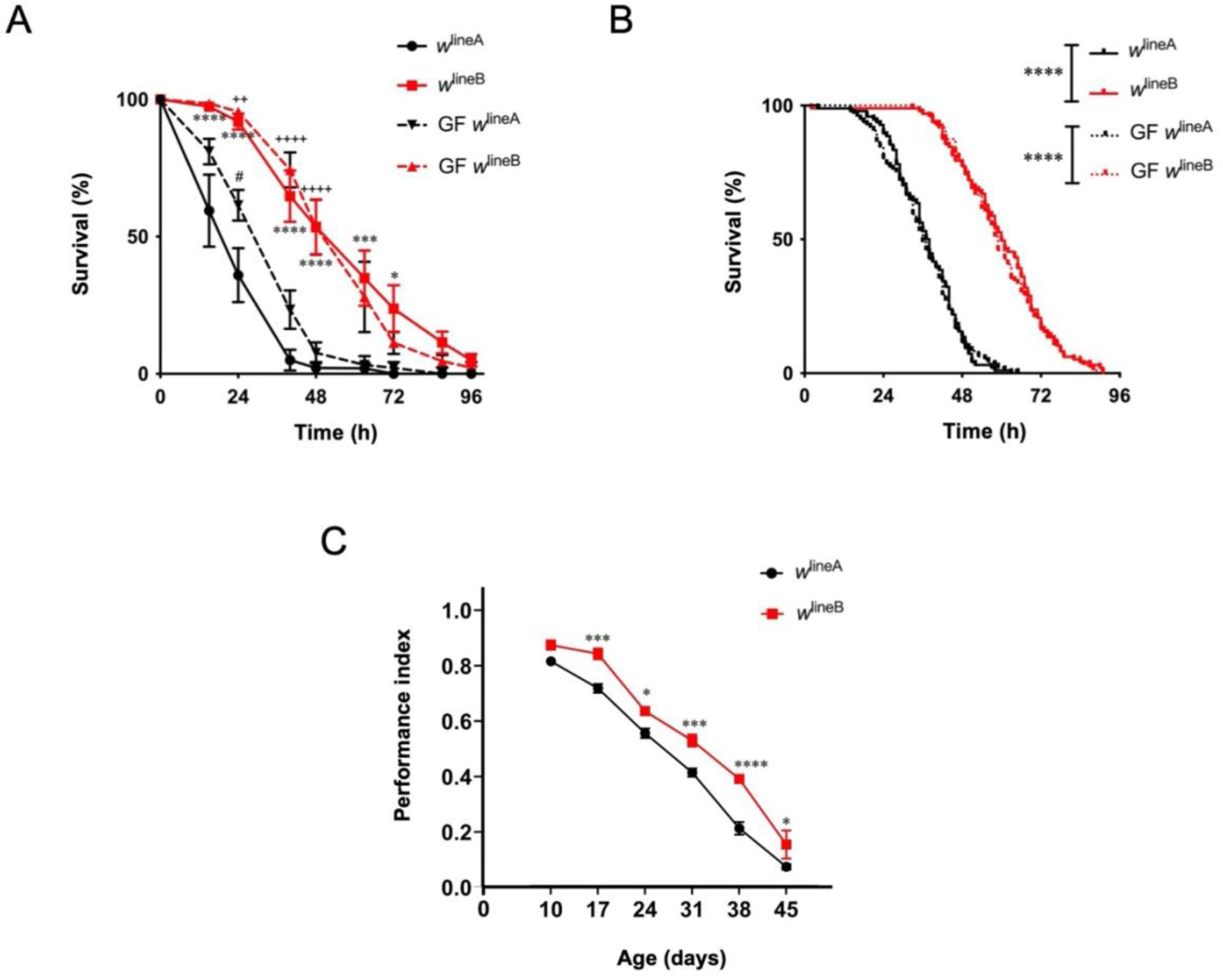
Differential stress resistance and locomotor performance of the two *Drosophila w*^1118^ lines. (**A**) Resistance to oxidative stress: *w*^lineA^ flies (*black curves*) were markedly more susceptible to oxidative stress than *w*^lineB^ flies (*red curves*), either when raised under standard (*solid curves*) or germ-free (GF) (*dashed curves*) conditions. Raising flies in GF condition slightly increased the resistance of *w*^lineA^ but did not change *w*^lineB^ resistance. Data represent percent survival as a function of time during continuous exposure to 20 mM paraquat. The results are mean of at least three independent experiments, each performed with about 100 flies per genotype or condition. Two-way ANOVA with Šídák’s multiple comparisons test (\*\*\*\**p* < 0.0001, \*\*\**p* < 0.001, \**p* < 0.05 for differences between the two *w*^1118^ lines in standard condition; ^++++^*p* < 0.0001,^++^*p* < 0.01 for differences between the two lines in GF condition; ^#^*p* < 0.05 is for differences between *w*^lineA^ in standard and GF conditions). (**B**) Resistance to starvation: *w*^lineA^ flies also showed a much lower resistance to nutrient deprivation than *w*^lineB^ (*solid curves*). Raising the flies in GF conditions (*dotted curves*) had no effect on the starvation resistance of either control line. Pooled data from three independent experiments, each carried out with 32 flies per genotype. Log-rank (Mantel-Cox) test (\*\*\*\**p* < 0.0001). (**C**) Evoked locomotion: from two weeks of age onwards, *w*^lineA^ flies (*black curve*) exhibited lower climbing performance in a startle-induced negative geotaxis (SING) test than the *w*^lineB^ line (*red curve*). Results are mean ± SEM of three independent experiments, each with 50 flies per genotype.

Recent reports indicate that the composition of the intestinal microbiota can modify resistance to oxidative stress or provide protection against nutritional stress in *Drosophila* (Consuegra et al., 2020; Onuma et al., 2023). This suggests that the difference in stress resistance of *w*^lineA^ and *w*^lineB^ could result from different gut microbiota. We therefore raised these flies in germ-free (GF) conditions to modulate the microbiome and analyzed consequences for susceptibility to oxidative and nutritional stress. Remarkably, the paraquat resistance of GF-raised *w*^lineA^ flies was found to be markedly increased: median survival time was 29 hours against 19 hours for conventionally reared (CR) *w*^lineA^ flies. In contrast, the paraquat resistance of *w*^lineB^ was not significantly different between GF and CR flies (Fig. 1A). Consequently, modulation of the microbiome mitigated the difference in oxidative stress resistance between *w*^lineB^ and *w*^lineA^ flies. This indicates that composition of the gut microbiota could be in part responsible for the difference in oxidative stress susceptibility between these two lines. However, we observed that raising flies in GF conditions did not affect the survival rate of either *w*^lineA^ or *w*^lineB^ under nutritional deprivation, as compared to CR flies (Fig. 1B). The gut microbiome, therefore, does not appear to be involved in the prominent difference in resistance to starvation between these two lines, leaving this question unresolved.

The gut microbiome has been reported to regulate locomotor behavior and aging in *Drosophila* (Schretter et al., 2018; Westfall et al., 2018; Lee et al., 2019; Arias-Rojas and Iatsenko, 2022). We then compared the age-related locomotor capacity of the two *w*^1118^ lines. The climbing reaction evoked by a gentle mechanic shock (startle) decreases with age in flies (Miquel et al., 1976; Grotewiel et al., 2005; White et al., 2010; Jones and Grotewiel, 2011; Riemensperger et al., 2011;Vaccaro et al., 2017) and it is one of the major indexes to evaluate the motor deficits in fly models of neurological disorders, such as PD (Feany and Bender, 2000; Riemensperger et al., 2013; Ordonez et al., 2018; Rahmani et al., 2022). Using startle-induced negative geotaxis (SING) to assess fly reactivity and climbing performance (Feany and Bender, 2000; Riemensperger et al., 2013), we found that the time course of age-related locomotor decline was similar in the two *w*^1118^ lines. However, *w*^lineB^ constantly exhibited better performance than *w*^lineA^ from middle age (17 days) to old age (45 days), suggesting that this line has a slower rate of locomotor aging than *w*^lineA^ (Fig. 1C). Such a difference in locomotor performance for individual flies could be related to the lower stress resistance of *w*^lineA^.

### Relative abundance of commensal bacteria species is divergent between the two control lines

The fact that microbiome modulation in germ-free conditions increased the oxidative stress resistance of *w*^lineA^, but not *w*^lineB^, prompted us to analyze their gut microbiome composition. For that, DNA from dissected whole gut of 10-day-old flies was prepared and sent for microbiome analysis by 16S ribosomal RNA (rRNA) gene sequencing, which is the most common approach in microbial classification (Sarangi et al., 2018). The effective reads generated after DNA amplification were 97,181 and 113,665 for *w*^lineA^ and *w*^lineB^, respectively, and the percentage of effective reads compared to raw reads was 88.40% for *w*^lineA^ and 80.19% for *w*^lineB^. No significant difference was observed for effective reads and percentage of effective reads between the two lines. The data were analyzed after excluding *Wolbachia*, because these bacteria are intracellular endosymbionts that are absent from the gut lumen. We observed that the most abundant order was *Acetobacterales* (34.3% and 52.2% for *w*^lineA^ and *w*^lineB^, respectively) without significant difference between the two fly lines. No significant differences were found either between for other bacterial orders, including *Corynebacteriales*, *Enterobacterales* and *Lactobacillales*. In contrast, *Clostridiales* were found to show higher relative abundance in *w*^lineA^ than in *w*^lineB^ (16.73% and 1.82%, respectively, *p* < 0.001), although both fly lines were maintained under identical culture conditions (Fig. 2A). We then analyzed *Wolbachia* and found that it was the most abundant genus in *w*^lineA^ (87.2%) and significantly less represented in *w*^lineB^ flies (2.6%) (*p* < 0.0001) (Fig. 2B). This suggests that the higher oxidative stress susceptibility of *w*^lineA^ could be caused by an imbalance of its intestinal microflora in favor of *Clostridiales* and *Wolbachia*.

**Figure 2.**
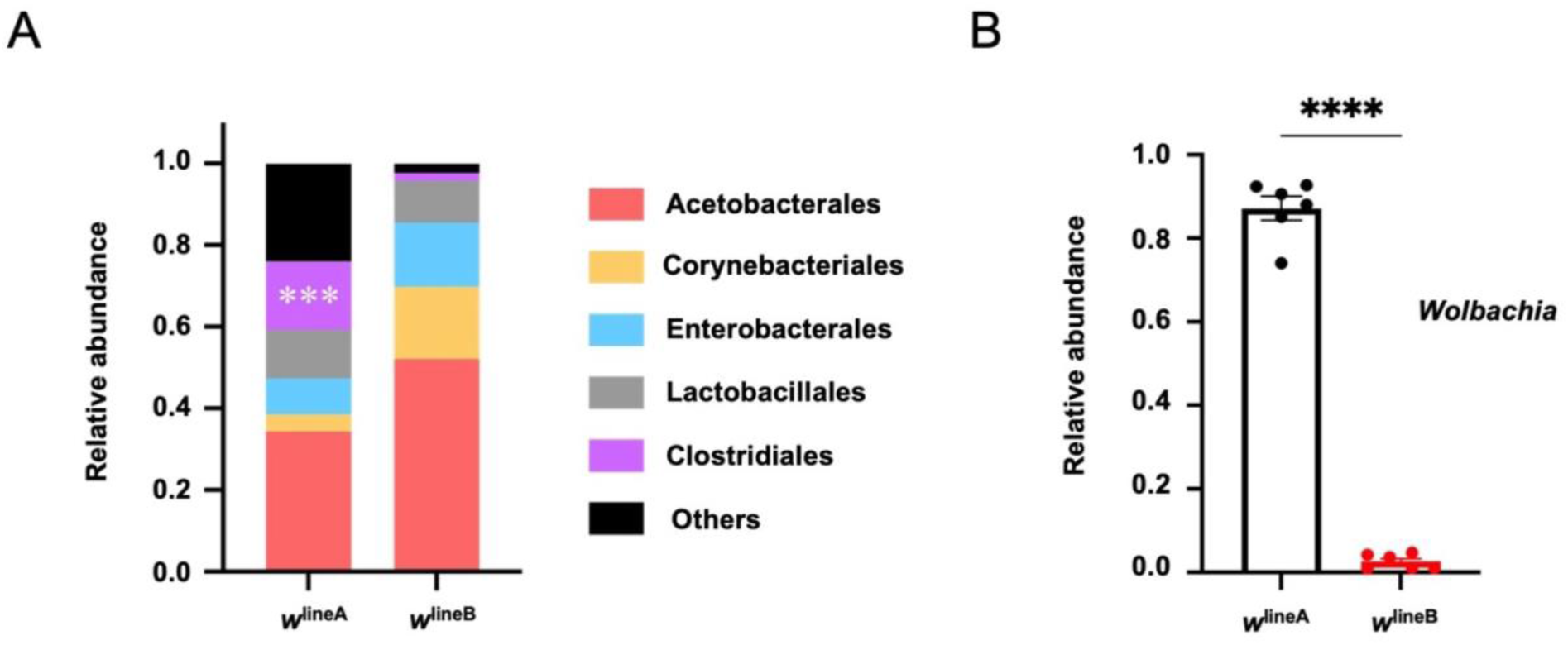
Divergent gut microbiota composition of the two *w*^1118^ lines. (**A**) Relative abundance of bacterial orders in the gut microbiome, excluding *Wolbachia*. *Acetobacterales* were predominant in both fly lines. *w*^lineA^ had higher abundance of *Clostridiales* than *w*^lineB^, while there were no significant differences for other orders. (**B**) Relative abundance of *Wolbachia* among bacterial genera in the gut microbiome. *Wolbachia* was much more abundant in *w*^lineA^ than in *w*^lineB^. Results are mean ± SEM of 6 replicates from two independent experiments, with 20 flies in each replicate and 120 flies in total per genotype. Data were analyzed with the Student’s *t*-test (\*\*\**p* < 0.001, \*\*\*\**p* < 0.0001).

### The two control lines have different levels of immune effectors but similar bacterial infection resistance

Immune effectors such as antimicrobial peptides (AMPs) and lysozymes contribute to the innate response in insects when challenged by various microorganisms. Seven families of AMPs have been identified in *Drosophila melanogaster* with antibacterial or antifungal activities according to AMP subtypes. They are mainly produced in the fat body and the gut, and strongly overexpressed under systemic infection (Imler and Bulet, 2005; Hanson and Lemaitre, 2020). Lysozymes are part of the system of inducible antibacterial defense but with a main function in the digestion of bacteria in *Drosophila*. About 17 genes encode putative lysozymes in the fly, and some of them are specifically expressed in the digestive tract (Kylsten et al., 1992; Daffre et al., 1994; Regel et al., 1998).

AMP expression is known to be involved in stress tolerance in *Drosophila* (Zhao et al., 2011) and to increase with age (Pletcher et al., 2002; Hanson and Lemaitre, 2020). More recently, it has also been reported that AMPs and lysozymes control the composition and abundance of the fly gut microbiota (Marra et al., 2021). The differences in stress resistance, aging and gut microbiome composition between *w*^lineA^ and *w*^lineB^ could thus in part reflect variations in expression levels of these immune effectors. We therefore quantified relative transcript levels of AMPs in the head and gut, and of lysozymes in the gut, for the two control lines. As shown in Fig. 3A, mRNAs of the Attacin-A, Defensin and Drosocin AMPs were found to be significantly less abundant in the heads of the less stress resistant *w*^lineA^ than in *w*^lineB^, whereas no difference was observed for Diptericin A and Drosomycin. Similarly in the gut, *w*^lineA^ also had significantly lower mRNA levels of the Attacin-A, Defensin and Drosocin, and also of the antibacterial Diptericin A and lysozymes, as compared to *w*^lineB^ (Fig. 3B). Like in heads, we found similar transcript levels of the antifungal Drosomycin in the gut between the two *w*^1118^ lines. The largest differences between the two lines were observed for Defensin, whose mRNA levels were 6.7 and 35.2-fold lower on average in the head and gut of *w*^lineA^, respectively, and for the lysozymes whose transcript levels were as much as 359-fold less abundant in the gut in *w*^lineA^ than in *w*^lineB^.

**Figure 3.**
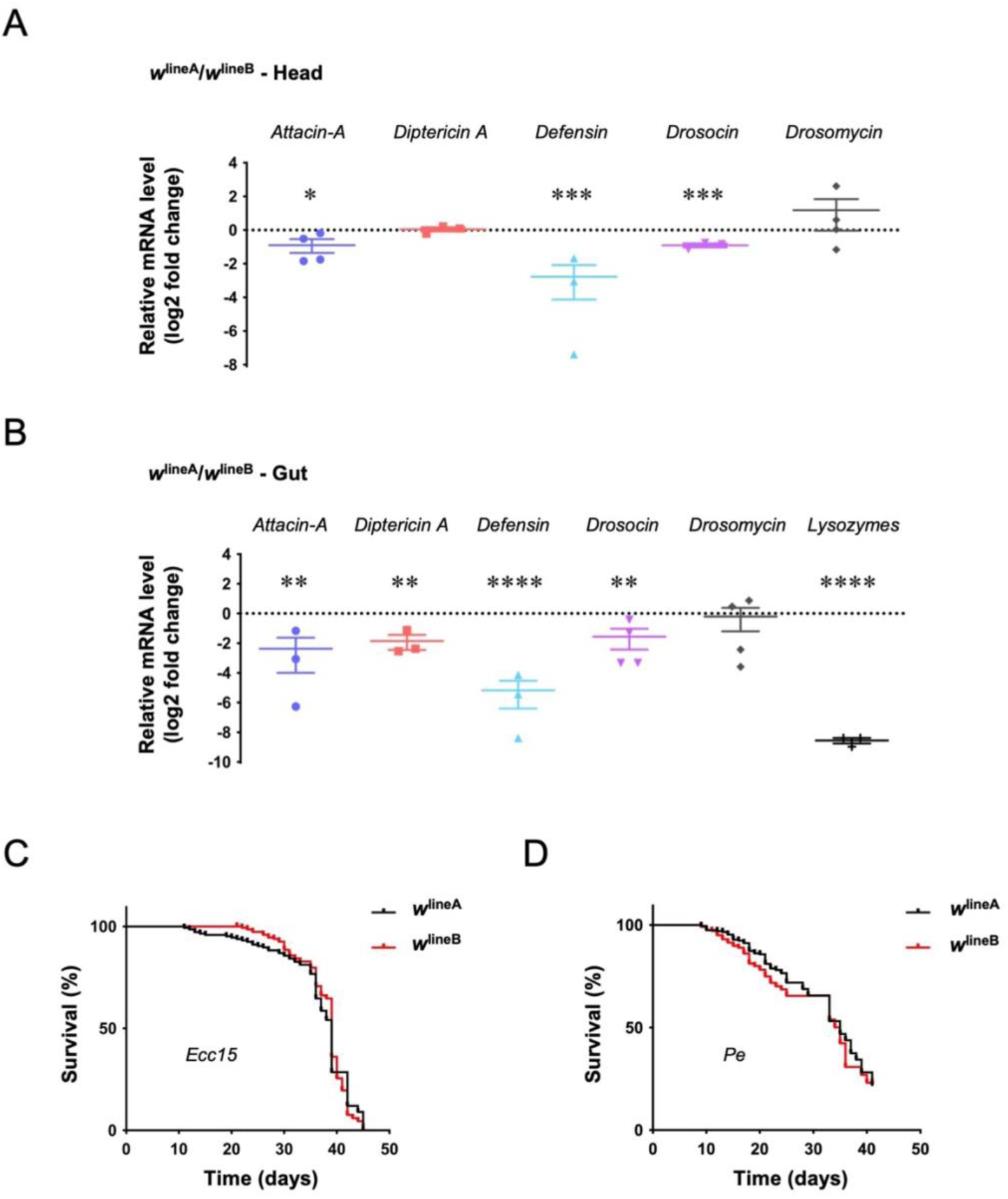
Transcript levels of immune effectors and infection resistance of the two *w*^1118^ lines. (**A**) In heads, mRNA levels of the AMPs *Attacin-A*, *Defensin* and *Drosocin* were found to be significantly lower in *w*^lineA^ than in *w*^lineB^. In contrast, no difference was found for *Diptericin A* and the antifungal peptide *Drosomycin*. (**B**) Comparable results were obtained in guts, with, additionally, lower levels of *Diptericin A* and lysozyme mRNAs in *w*^lineA^ than in *w*^lineB^. Results in **A** and **B** are mean ± SEM of at least three independent experiments, each carried out with twenty 7-day-old adult flies per genotype. Student’s *t*-test (\*\*\*\**p* < 0.0001 \*\*\**p* < 0.001, \*\**p* < 0.01 and \**p* < 0.05). (**C**, **D**) Survival curve of *w*^lineA^ and *w*^lineB^ following oral infection with *Erwinia carotovora carotovora 15* (*Ecc15*) (**C**) and *Pseudomonas entomophila* (*Pe*) (**D**). *w*^lineA^ and *w*^lineB^ flies showed similar susceptibility to pathogen bacterial infection. Data in **C** and **D** were pooled from 2 or 3 independent experiments with 50 flies in each group and analyzed by the log-rank (Mantel-Cox) test.

The lower levels of several immune effectors in the head and gut of *w*^lineA^ could potentially compromise the resistance of these flies to infection by pathogenic bacteria. We then compared the survival of the two *w*^1118^ lines after the oral ingestion of two bacterial pathogens, *Erwinia carotovora carotovora 15* (*Ecc15*) and *Pseudomonas entomophila* (*Pe*). *Ecc15* is a phytopathogen that uses flies as vectors and is normally nonlethal to *Drosophila*, although it increases AMP expression in its host and triggers intestinal tissue damage and a potent avoidance behavior (Basset et al., 2000; Charroux et al., 2020; Lei et al., 2022). *Pe* is highly pathogenic for insects and its ingestion induces AMP expression and irreversible gut damage (Vodovar et al., 2005). It has been reported that *Drosophila* lines with better resistance to oxidative stress and paraquat were also more resistant to *Pe* infection (Bou Sleiman et al., 2015). However, we observed that *w*^lineA^ and *w*^lineB^ flies unexpectedly have a similar lifespan after oral ingestion of these bacteria when cultured on an otherwise sterile medium. Their median lifespans were about 39 days when infected with *Ecc15* (Fig. 3C) and 35 days with *Pe* (Fig. 3D). These results suggest that the difference in basal levels of AMPs and lysozymes in the two *w*^1118^ lines does not compromise their immune response in the case of a systemic infection.

### Age-related locomotor defects induced by pan-neuronal α-synA30P depend on background of the transgenic flies

The fact that the background of the treated lines had a major influence on fly survival in the drug-induced paraquat model of PD (Fig. 1A) led us to investigate whether it could also affect motor symptoms in a transgenic model of this disease. Expression of the pathogenic A30P mutant form of human α-synuclein (α-synA30P) in *Drosophila* neurons triggers age-related locomotor defects (Feany and Bender, 2000; Riemensperger et al., 2013; Chen et al., 2014; Issa et al., 2018; Rahmani et al., 2022). Accordingly, as shown in Fig. 4A, expressing α-synA30P in all neurons with the *nSyb-Gal4* driver using non-backcrossed parent lines robustly led to climbing impairments that were statistically significant starting from 17 days after adult eclosion (Fig. 4A, *yellow curve*). However, after the parent *nSyb-Gal4* driver and *UAS-α-synA30P* effector lines were backcrossed for seven generations in the stress-resistant *w*^lineB^ background, the locomotor defects induced by α-synuclein were found to be delayed and considerably mitigated, with impairments not noticeable before 24 days after eclosion. While climbing performance was nearly abrogated at 31 days in non-backcrossed flies expressing *α-synA30P* pan-neuronally (Fig. 4A, *yellow curve*), it was maintained and only moderately reduced at the same age in comparison to controls in the *w*^lineB^ background (Fig. 4A, *brown curve*). The background of the transgenic line can therefore significantly alter motor impairments induced by pan-neuronal α-synuclein expression in *Drosophila*, which could contribute to the variability of results observed with this PD model in different laboratories.

**Figure 4.**
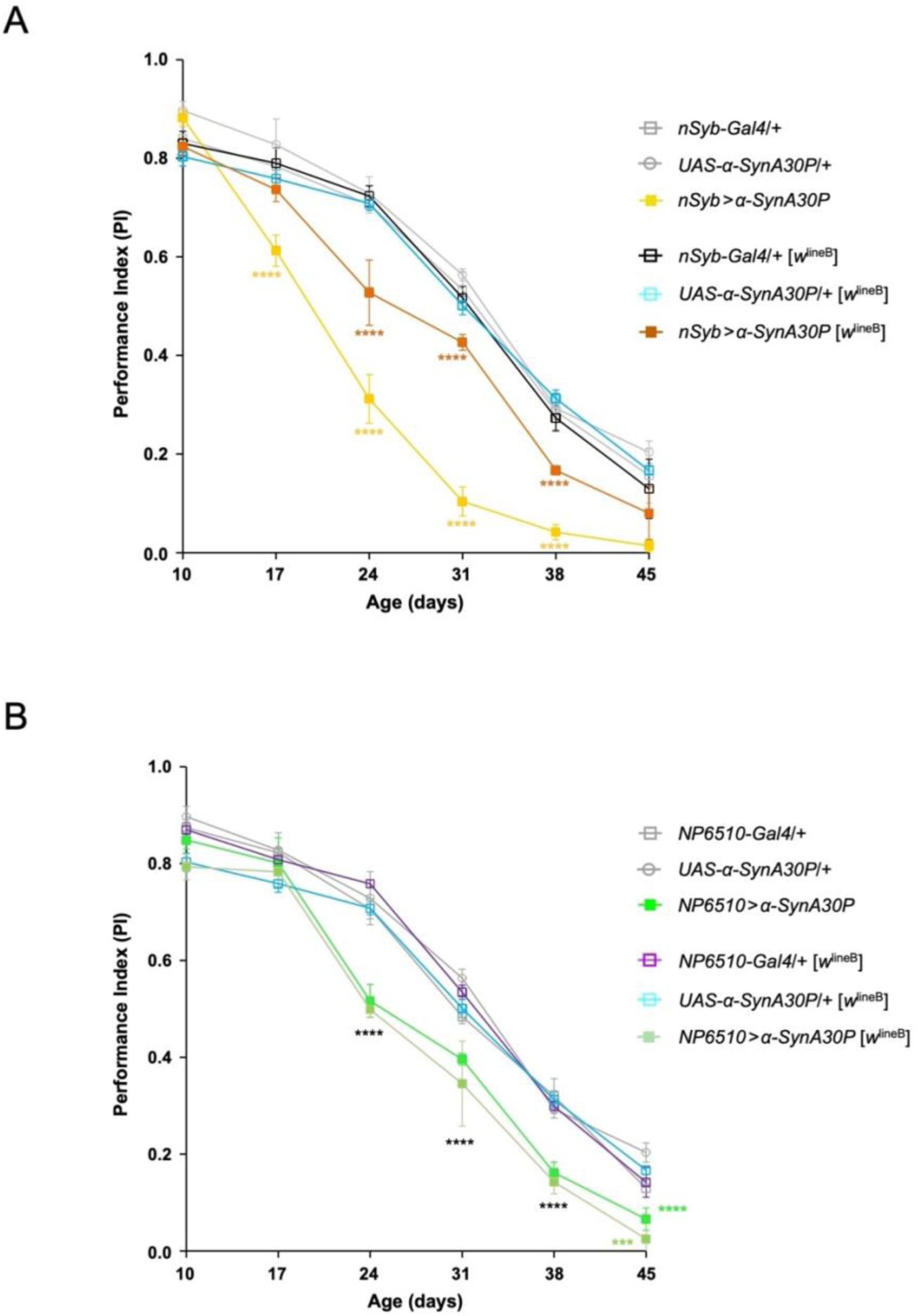
Influence of the transgenic line background on age-dependent locomotor defects induced by mutant α-synuclein. (**A**) Using non-backcrossed parent lines, the pan-neuronal expression of mutant α-synuclein (α-synA30P) with the driver *nSyb-Gal4* induced prominent startle-induced climbing defects that started from 17 days of age (*yellow curve*), as compared to heterozygous driver and *UAS-α-synA30P* alone controls (*grey curves*). In contrast, climbing defects were strongly attenuated using driver and UAS parent lines backcrossed in the stress-resistant *w*^lineB^ background (*brown curve*), without altering SING performance of the driver and UAS alone controls (*black and blue curves*), which remained similar at all ages to that of non-backcrossed controls (*grey curves*). (**B**) In contrast, using either non-backcrossed parent lines or after their backcross into the *w*^lineB^ background, selective α-synA30P expression in a subset of the protocerebral anterior medial (PAM) dopaminergic neurons with the driver *NP6510-Gal4* triggered similar SING defects (*green curves*) compared to respective driver and UAS effector alone controls (*grey, purple and blue curves).* In all panels, results are mean ± SEM of three independent experiments, each with 50 flies per genotype. Two-way ANOVA with Tukey’s post-hoc test (\*\*\*\**p* < 0.0001, *** *p* < 0.001, \**p* < 0.05).

Locomotor impairments can also be induced in flies by selectively expressing α-synA30P in a subset of 15 dopaminergic neurons of the brain protocerebral anterior medial (PAM) cluster, using the driver *NP6510-Gal4* (Riemensperger et al., 2013; Rahmani et al., 2022; Kajtor et al., 2025). In contrast to pan-neuronal expression, we observed similar age-related climbing impairments in transgenic flies expressing α-synA30P selectively in these PAM neurons, whether the driver and *UAS* parent lines were non-backcrossed or backcrossed in the *w*^lineB^ background (Fig. 4B, *green curves*). This finding intriguingly suggests that the vulnerability of these PAM dopaminergic neurons to α-synA30P neurotoxicity is less influenced than for other brain neurons by the general background of the fly.

## Discussion

*Drosophila* models of human neurological diseases generally rely on comparisons between healthy and diseased flies, the latter having for instance defective stress resistance or accelerated age-related locomotor decline. However, the distinction between the normal physiological state and pathological states is often difficult to make. In a famous epistemological study, the philosopher Georges Canguilhem challenged the idea that the pathological state is merely a quantitative modification of the physiological state and argued that the pathological state can also be considered as normal, since “the morbid state is always a certain mode of living” (Canguilhem, 1991). Here we find that two independent *Drosophila* lines of the same *w*^1118^ strain, which is commonly used as healthy normal flies in disease model studies, exhibit dramatic differences in their susceptibility to nutrient starvation and oxidative stress. We report that differences in gut microbiome composition and relative expression of immune effectors can play a prominent role in such variability. We also show that the locomotor impairments induced by mutant α-synuclein in a transgenic Parkinson disease model can be significantly alleviated in a strongly stress-resistant background. This indicates that the same pathological conditions could lead to variable, and sometimes opposite, conclusions, depending on the control line that is used for comparison. The validity and reproducibility of a *Drosophila* model, therefore, would largely depend on a careful selection and previous characterization of the reference line.

### Role of the gut microbiota in stress resistance of the reference lines

It has been recently reported that the composition of the intestinal microbiota can modify resistance to oxidative stress or provide protection against nutritional stress in *Drosophila* (Consuegra et al., 2020; Onuma et al., 2023). Accordingly, we found that raising flies in germ-free conditions had no effect on survival of the stress-resistant *w*^lineB^ flies exposed to paraquat, but increased significantly that of the less resistant *w*^lineA^. In contrast, germ-free culture conditions did not alter the susceptibility of the two control lines to nutrient starvation, suggesting that the high difference in starvation resistance between the two *w*^1118^ lines likely originates from the genomic background rather than the gut microbiota.

Analysis of the gut microbiome composition by 16S rRNA sequencing showed that *w*^lineA^ has significantly higher abundance of *Clostridiales* and intracellular *Wolbachia* than *w*^lineB^. These bacteria could undermine the oxidative stress resistance of *w*^lineA^. The strictly anaerobic *Clostridiales* is a dominant bacterial order in mammals, which is often minor but sometimes of higher relative abundance in the *Drosophila* gut (Chandler et al., 2011; Walters et al., 2020). The physiological impact of these bacteria in insects has not been specifically studied. *Wolbachia* is a genus of intracellular gram-negative bacteria found in arthropods and nematodes, with parasitic or symbiotic effects according to the host species (Werren et al., 2008). The presence of this endosymbiont was reported to modify the composition of gut microbial species in *Drosophila* (Simhadri et al., 2017). This could be a reason why *Clostridiales* are more abundant in *w*^lineA^ than in *w*^lineB^, in addition to host genetic variation. Moreover, the production of reactive oxygen species (ROS) is significantly increased in *Wolbachia*-infected mosquito cells in culture compared to cured cells (Brennan et al., 2008) and in *Drosophila* hosting viral-protective *Wolbachia* strains (Wong et al., 2015). *Wolbachia* can also increase host sensitivity to paraquat in insects, as shown in the parasitic wasp *Asobara japonica* (Monnin et al., 2017) and in *Drosophila* where it depended on the fly genetic background (Capobianco III et al., 2018). Besides, *Wolbachia* is known to be potently antiviral (Hedges et al., 2008; Teixeira et al., 2008; Bian et al., 2010) and can also protect against fungus pathogens in *Drosophila* (Perlmutter et al., 2025). It is noteworthy that around 30% of the fly lines available in the Bloomington *Drosophila* Stock Center are colonized by *Wolbachia* (Clark et al., 2005), as is the case for a large proportion of laboratory strains (Min and Benzer, 1997).

### Variations in basal immune effector levels could underlie differences between the two control lines

In addition to their role in the innate immune response against pathogen infection, AMPs have been reported to be involved in stress resistance, neurodegeneration and microbiome dysbiosis in *Drosophila* (Zhao et al., 2011; Cao et al., 2013; Shukla et al., 2019; Hanson and Lemaitre, 2023). AMPs and lysozymes also control the structure and diversity of the fly gut microbial community (Marra et al., 2021). Variations in their expression levels could therefore contribute to the differences between *w*^lineA^ and *w*^lineB^. We observed that the stress-resistant line *w*^lineB^ generally had higher relative transcript levels of these immune effectors in the gut and head. It was the case for Attacin-A, Defensin, Drosocin, Diptericin A and lysozymes in gut, and for Attacin-A, Defensin and Drosocin in head, with the exception of the antifungal Drosomycin that did not vary between *w*^lineA^ and *w*^lineB^. Attacin-A, Drosocin and Diptericin A are active against gram-negative bacteria, while Defensin, which is the AMP that differs the most between the two lines in both head and gut, is active against gram-positive bacteria. Lysozyme levels were also found to be very much decreased in the intestine of *w*^lineA^ flies.

The fact *w*^lineA^ has both enrichment in *Wolbachia* and low AMP levels in gut is not contradictory, as *Wolbachia* do not induce AMP synthesis in insects (Bourtzis et al., 2000). In addition to its pro-oxidant effect, paraquat exposure also strongly downregulates AMP expression in *Drosophila* (Maitra et al., 2019), while conversely starvation was shown to increases AMP expression in fly larvae (Becker et al., 2010). Higher levels of the AMP Diptericin have been correlated to a better resistance to hyperoxia-induced oxidative stress through increased antioxidant enzyme activities (Zhao et al., 2011). Overall, this suggests that the remarkable tolerance of *w*^lineB^ to starvation and oxidative stress could be related to its higher basal levels of immune effectors.

In contrast, we observed that the survival rate of *w*^lineA^ and *w*^lineB^ was similar after ingestion of two pathogenic bacteria, *Ecc15* and *Pe*, that induce intestinal tissue damage (Basset et al., 2000; Vodovar et al., 2005). The differences in microbiome composition and basal immune effector levels appear therefore not sufficient to alter their immune response against systemic bacterial infection.

### Background of the transgenic line significantly affects α-synuclein-induced motor defects

Startle-induced climbing responses decline progressively with age in wild-type flies and are abolished at about 50 days at 25°C, approximately two weeks before *Drosophila* maximal lifespan (Miquel et al., 1976; Grotewiel et al., 2005; White et al., 2010; Jones and Grotewiel, 2011; Riemensperger et al., 2011; Vaccaro et al., 2017). In contrast, spontaneous locomotion does not vary during adult life and even increases in old flies (White et al., 2010). It has been reported that the gut microbiome can regulate spontaneous locomotion in *Drosophila* (Schretter et al., 2018). We have indeed observed that the stress-resistant *w*^lineB^ performed better than *w*^lineA^ in startle-induced climbing from middle age onwards. Nevertheless, compared to the dramatic differences observed with oxidative stress susceptibility and starvation resistance, the difference in climbing performance between *w*^lineA^ and *w*^lineB^ appeared relatively moderate.

The expression of human wild-type or mutant α-synuclein in *Drosophila* neurons has been widely shown to induce age-related locomotor impairments and loss of brain dopaminergic neurons (Feany and Bender, 2000; Barone et al., 2011; Riemensperger et al., 2013; Chen et al., 2014; Wang et al., 2015; Issa et al., 2018; Mohite et al., 2018; Ordonez et al., 2018; Bridi et al., 2021; Rahmani et al., 2022). However, these effects have been found to be variable in different laboratories or sometimes difficult to reproduce (Pesah et al., 2005; Navarro et al., 2014). Here we found that backcrossing the transgenic lines in the *w*^lineB^ background significantly delayed and much alleviated the negative geotaxis defects triggered by pan-neuronal expression of α-synA30P, compared to results with non-backcrossed lines. The use of a less sensitive climbing assay could have led us to conclude that α-synA30P did not trigger any locomotor impairments in these conditions. This shows that variations in α-synuclein-induced defects could partly result from the background of the transgenic lines in which the pathogenic protein is expressed.

Locomotor defects can also be observed when α-synuclein expression is restricted to a subset of 15 PAM dopaminergic neurons with the NP6510-Gal4 driver (Riemensperger et al., 2013; Rahmani et al., 2022; Kajtor et al., 2025). We observed that the climbing impairments were not mitigated in the *w*^lineB^ background in this case, in contrast to pan-neuronal α-synA30P expression. The vulnerability of these PAM neurons towards α-synuclein seems to be less sensitive to the fly line background for unknown reasons. Interestingly, the locomotor defects induced by α-synA30P expression in all neurons in the *w*^lineB^ background (Fig. 2A, *red curve*) and in PAM neurons using both non-backcrossed or backcrossed parent lines (Fig. 2B, *green curves*) look quantitatively very similar, with climbing impairments starting from 24 days in both cases and the same level of reduction in motor performance at all later ages. It could then be hypothesized that there are two components in the locomotor disturbance induced by pan-neuronal α-synuclein expression in flies: a component resulting from α-synA30P expression in the PAM dopaminergic neurons, which would be maintained in the *w*^lineB^ background, and a component resulting from α-synA30P expression in other neurons that would in contrast be strongly mitigated in a stress-resistant background.

## Conclusion

Overall, our results show that two apparently identical *Drosophila* control lines can exhibit highly significant differences in their susceptibility to various stress and disease-causing agents. The causes of these differences can be quite intricate, involving a combination of intestinal microbiome species diversity, divergent immune effectors levels, silent genomic variations and possibly other factors. Such effects could partly explain the difficulties sometimes encountered in reproducing what was previously observed in neurodegeneration models. Taking in account several characteristics of the reference line, and not only the visible part of its genotype, would therefore be useful to improve the consistency of disease modeling studies in *Drosophila*.

## Materials and Methods

### *Drosophila* maintenance and strains

Conventionally reared (CR) flies were maintained at 25°C on standard cornmeal-yeast-agar medium supplemented with the antifungal agent methyl-4-hydroxybenzoate, under a 12h:12h light-dark cycle. Two separate *w*^1118^ lines were used in this work, named *w*^lineA^ and *w*^lineB^, respectively. *w*^lineA^ was obtained from the Bloomington *Drosophila* Stock Center (RRID:BDSC_3605) several years ago and *w*^lineB^ was received from BestGene (Chino Hills, CA, USA) more recently. The other fly strains used were: *nSyb-Gal4* from BDSC (RRID:BDSC_51635), *NP6510-Gal4* from the Kyoto *Drosophila* Stock Center (DGRC_113956), and *UAS-α-synA30P*, also named *UAS-Hsap\SNCA.A30P* (generous gift from Mel Feany, Harvard Medical School, Boston, USA). Prior to some experiments, *nSyb-Gal4*, *NP6510-Gal4* and *UAS-α-synA30P* were backcrossed for seven generations with the *w*^lineB^ line.

### *Drosophila* culture in axenic conditions

*Drosophila* culture in germ-free (GF) conditions was carried out as previously described (Téfit and Leulier, 2017) with minor modifications. Briefly, non-virgin females (5 to 7 day-old) were collected and kept in glass tubes on standard cornmeal-yeast-agar medium for 2 days. They were then transferred to egg-laying chambers composed of an empty bottle with holes on the wall for air circulation covered by a 60 mm-diameter plate containing grape juice-based agar medium and a drop of yeast paste. Eggs were collected after 20 hours with a brush, transferred to a nylon basket and successively washed for 2 min in baths of 2.7% bleach, 70% ethanol and sterile water. Bleached eggs were deposited on autoclaved standard culture medium supplemented with an antibiotic cocktail containing 50 μg ampicillin sodium salt (Euromedex, EU0400), 50 μg kanamycin monosulfate (Euromedex, UK0010), 50 μg tetracycline hydrochloride (Euromedex, UT2965) and 15 μg erythromycin (Euromedex, TO-E002) per liter of fly food. Emerging adult females were maintained on antibiotic-supplemented standard medium and tested for axenicity by incubating whole-body lysates on Luria broth-agar (Carl Roth, X969), MRS-agar (Carl Roth, X924) and Mannitol salt-agar (Carl Roth, CL81.1) plates.

### Startle-induced locomotion assay

Evoked climbing activity was monitored by a startle-induced negative geotaxis (SING) test as previously described (Riemensperger et al., 2013). 50 adult males divided into 5 groups of 10 flies were placed in vertical pipettes (25-cm long, 1.5-cm diameter, Greiner Bio-One, 760180) and left for about 30 min for habituation. Then, columns were tested individually by gently tapping them down (startle), to which flies normally responded by climbing up. After 1 min, flies having reached at least once the top of the column (above 22 cm) and flies that never left the bottom (below 4 cm) were counted. Each fly group was assayed three times at 15 min intervals. The performance index (PI) for each column is defined as ½[1 + (n_top_-n_bot_)/n_tot_], where n_tot_ is the total number of flies in the column, n_top_ the number of flies at the top and n_bot_ the number of flies at the bottom after 1 min. Age-related SING performances were tested weekly over 6 weeks, starting on day 10 after adult eclosion. Three independent experiments were carried out per genotype.

### Oxidative stress and starvation resistance

*Drosophila* oxidative stress resistance was measured by dietary ingestion of the herbicide paraquat (methyl viologen dichloride hydrate; Sigma-Aldrich, 856177) as previously described (Cassar et al., 2015) with minor modifications. Non-virgin 7-day-old females (100 flies per genotype) were starved for 2 hours in empty vials at 25°C, then quickly anesthetized on ice and transferred to 35 mm diameter Petri dishes (Dutscher, 353001) (10 flies per dish) containing two layers of Whatman blotting paper (Sigma-Aldrich, WHA3001917) soaked with 400 µl of 20 mM paraquat in 2% (w/v) sucrose (Euromedex, 200-301). The Petri dishes were then placed in a plastic box under saturated humidity conditions and incubated at 25°C. Fly survival was scored at several time points over 4 days. Experiments were carried out three times independently with 300 flies per genotype in total and results at a given time point correspond to the mean of the scores obtained in each independent experiment.

Resistance to nutrient starvation was determined using DAM2 *Drosophila* Activity Monitors (Trikinetics). 7-day-old females (32 flies per genotype) were placed individually in Trikinetics transparent glass tubes containing 1.5% agar in sterile water and incubated at 25°C up to 5 days under a 12h:12h light-dark cycle. Flies that stopped moving back and forth for 30 min and did not move again were considered dead. Results are presented as percent survival as a function of time.

### Gut DNA sample preparation

DNA samples were prepared from 10-day-old non-virgin females. In each experiment, three lots of 20 *w*^lineA^ or *w*^lineB^ flies were placed in sterile 1.5ml Eppendorf tubes and washed twice with 70% ethanol and once with sterile water for 1 min by a gentle vortex. Whole guts were dissected and immediately processed for DNA extraction using a protocol adapted from the genomic DNA prep of E. Jay Rehm (Berkeley *Drosophila* Genome Project) (Bellen et al., 2004) (https://www.fruitfly.org/about/methods/inverse.pcr.html). Briefly, 20 guts were homogenized for 100 s at maximal speed in 400 µl DNA extraction buffer (100 mM Tris-HCl, 100 mM NaCl, 100 mM EDTA, 0.5% (w/v) SDS, pH 7.5) using bead tubes in a Minilys apparatus (Bertin Instruments, Montigny-le-Bretonneux, France). Samples were incubated for 30 min at 65°C, then 800 µl of 4.3 M LiCl/1.4 M KAc solution was added and the samples were cooled on ice for 15 min. After 15 min centrifugation at 13,000 rpm (10,700 g), 1 ml of the supernatant was transferred into a new microcentrifuge tube, avoiding the floating crud, and 600 µl isopropanol was added and mixed by inverting. Precipitates were spun down for 17 min at 13,000 rpm, and the supernatants were discarded. The pellets were washed with 1 ml 70% ethanol and spun down again for 10 min at 13,000 rpm. After removal of the supernatant, the DNA pellet was air dried as much as possible, resuspended in 50 µl sterile water and stored at −20°C. DNA concentration was measured in a NanoDrop spectrophotometer and diluted to 1 µg/μL in sterile water. DNA purity was then monitored on 1% agarose gels.

### DNA sequencing and analysis

The gut DNA samples were sent to Novogene (Cambridge, UK) for sequencing and microbiome composition analysis. DNA was first amplified with the primers 515F and 907R with a barcode (515F: 5’-GTGCCAGCMGCCGCGGTAA-3’, 907R: 5’-CCGTCAATTCCTTTGAGTTT3’), which target the V4 and V5 region of the bacterial 16s rDNA. Sequencing was conducted on high-throughput Illumina sequencing platform which can realize the rapid-mode paired-end 250bp (PE250) sequencing strategy. Sequences with ≥97% similarity were assigned to the same Operational Taxonomic Units (OTUs). 6 DNA samples from independent extractions of *w*^lineA^ and *w*^lineB^ flies were sequenced in total.

### RNA extraction

RNAs were prepared from 20 heads or dissected guts from 10-day-old non-virgin females. Tissues were lysed and homogenized in 400 μL or 1.2 ml QIAzol Lysis reagent (Qiagen, 79306), respectively, for 100 s at maximal speed in the Minilys apparatus. Total RNA was extracted essentially as described (Green and Sambrook, 2020), except that the final precipitates were washed twice in 75% ethanol. RNA concentration was measured in a NanoDrop spectrophotometer and diluted to 1 µg/μL in RNAse-free water.

### Quantitative reverse transcription-coupled PCR (RT-qPCR)

1 µg RNA was reverse transcribed to cDNAs by using the Maxima First Strand cDNA Synthesis Kit (Thermo Fisher Scientific, K1671) with oligo(dT)20 primers (Thermo Fisher Scientific, 18418020). qPCR was carried out using a LightCycler 480 and the SYBR Green I Master mix (Roche LifeScience). The housekeeping gene *RpL32* (*rp49*) was used as internal control. The program cycles were: 10 s denaturation at 95°C, 10 s annealing at 55°C or 60°C and 20 s elongation at 72°C, with 45 cycles in total. All reactions were performed in triplicate. Melting curves were analyzed to check the specificity of amplification products and relative mRNA quantification was carried out by the ΔΔCt method. Sequences of the forward and reverse primers used were: for *RpL32*, 5’-GACGCTTCAAGGGACAGTATC and 5’-AAACGCGGTTCTGCATGAG, for *Attacin-A* (*AttA*), 5’-ATGCAGAACACAAGCATC and 5’-CAGTTGTGACTGGACCACT, for *Diptericin A* (*DptA*), 5’-ACCATTGCCGTCGCCTTAC and 5’-CTCCATTCAGTCCAATCTCGTGG, for *Defensin* (*Def*), 5’-TCGTTCTCGTGGCTATCGC and 5’-TGAACCCCTTGGCAATGCA, for *Drosocin* (*Dro*), 5’-CGTTTTCCTGCTGCTTGCTT and 5’-GGCAGCTTGAGTCAGGTGAT, for *Drosomycin* (*Drs*), 5’-AGTACTTGTTCGCCCTCTTCG and 5’-GTATCTTCCGGACAGGCAGT, and for lysozymes, 5’-CTACAACGGCTCCAACGACT and 5’-GATGTCGTCGGTCAAGAGGG. The lysozyme primers amplified *Lysozyme B* (*LysB*), *LysD* and *LysE* at their consensus sequence.

### Infection assay

Survival assays following oral infection with *Erwinia carotovora carotovora 15* (*Ecc15*) and *Pseudomonas entomophila* (*Pe*) were conducted essentially as described (Liehl et al., 2006). *Ecc15* (generous gift from Julien Royet, Aix-Marseille Université, France) and rifampicin-resistant *Pe* mutants (generous gift from Bruno Lemaitre, École Polytechnique Fédérale de Lausanne, Switzerland) were incubated overnight in Luria broth medium with or without 100μg/ml rifampicin (Euromedex, 1059) at 30°C with agitation at 200 rpm. *Ecc15* or *Pe* was collected by centrifugation at 4,000 rpm for 15 min in 50-ml tubes and diluted with autoclaved 5% (w/v) sucrose to obtain a solution at optical density of 100. 50 female flies aged 7 days were starved for 2 hours at 25°C in empty vials and then transferred into a bottle containing sterile cornmeal-yeast-agar medium covered by 1 layer of pre-autoclaved Whatman blotting papers (Sigma-Aldrich, WHA3001917) soaked in 600 μL of *Ecc15* or *Pe* solution. The infection procedure was repeated at 3-day intervals.

### Statistical analysis

Statistical analyses were performed by the GraphPad Prism 6 software. Results of SING and paraquat resistance experiments were analyzed by two-way ANOVA with Šídák’s *post-hoc* pairwise comparisons test. Survival curves for starvation and infection experiments were generated and analyzed by the Log-rank test. Student’s *t*-test was used for comparison between two groups. Significant values in all figures: \**p* < 0.05, \*\**p* < 0.01, \*\*\**p* < 0.001, \*\*\*\**p* < 0.0001. Error bars in figures represent standard error of the mean (SEM).

## Acknowledgements

We would like to thank François Leulier for his advice on the procedure for rearing flies in germ-free conditions, Mel Feany for the gift of the *UAS-α-synA30P* line, and Baya Chérif-Zahar for helpful discussions. We also thank Bruno Lemaitre and Julien Royet for kindly providing the *Ecc15* and *Pe* bacterial strains, respectively, together with useful methodological suggestions for the infection assay.

## Author contributions

Conceptualization: X.Y., A.D., Z.R., and S.B.; Formal analysis: X.Y., A.D., A.H. and Z.R.; Investigation: X.Y., A.D., A.H. and Z.R.; Methodology: X.Y., A.D. and Z.R.; Writing–-original draft: X.Y. and S.B.; Writing–-review & editing: X.Y., A.D., Z.R., and S.B.; Supervision: S.B.; Project administration: S.B.; Funding acquisition: S.B.

## Data availability

All relevant data are included within the article. Raw data are available upon request.

